# Periostin exon 17 skipping enhances the efficacy of local AAV-microdystrophin administration in a fibrotic model of Duchenne muscular dystrophy

**DOI:** 10.1101/2025.01.21.633639

**Authors:** Jessica Trundle, Alexis Boulinguiez, Ngoc Lu-Nguyen, James March, Alberto Malerba, Linda Popplewell

## Abstract

Duchenne muscular dystrophy (DMD) is a severe, progressive genetic disorder primarily affecting boys, characterized by muscle degeneration due to mutations in the DMD gene encoding dystrophin, a crucial protein for muscle fiber integrity. The disease leads to significant muscle weakness and eventually to loss of ambulation. AAV-microdystrophin gene therapy shows promise in preclinical and clinical settings. However, muscle fibrosis, a consequence of chronic inflammation and extracellular matrix (ECM) remodeling, exacerbates disease progression and may hinder therapeutic efficacy. Periostin, a matricellular protein involved in fibrosis, is upregulated in DMD rodent models and correlates with collagen deposition. We previously developed an antisense oligonucleotide strategy to induce exon 17 skipping and so reduce periostin expression and collagen accumulation in the fibrotic D2.*mdx* mouse model of DMD. Here, we investigated the combined effects of periostin modulation and AAV-microdystrophin (AAV-MD1) treatment. We found that systemic periostin splicing modulation with intramuscular AAV-MD1 administration significantly improved muscle function, assessed by grip strength and treadmill performance, compared to either single treatment. Importantly, periostin exon skipping increases the microdystrophin protein expression. These findings suggest that targeting periostin in conjunction with microdystrophin therapy could represent a valid therapeutic strategy for DMD.

## Introduction

Duchenne muscular dystrophy (DMD) is a severe, progressive genetic disorder characterized by muscle degeneration and weakness. It affects approximately 1: 5,000 live male births worldwide^1^ and is caused by mutations in the DMD gene, which encodes dystrophin, a crucial protein for muscle fiber integrity^2^. Loss of ambulation occurs by the early teenage years and, without intervention, cardiac^3^ or respiratory failure^4^ cause death in early adulthood.

AAV-microdystrophin (AAV-MD) gene therapy stands as one of the most promising avenues for treating DMD. We developed AAV8-Spc5-12-MD1, a viral vector presently undergoing clinical trials (GNT0004)^5,6^ that expresses microdystrophin-1 (MD1). This shorter version of dystrophin protein showcases robust protein expression in murine and canine DMD models when delivered through Adeno-associated viral (AAV) vectors^7–11^. An AAV-microdystrophin clinical product has received FDA approval and is now available in the market (Elevidys®)^12^. While promising, published clinical data suggest that this approach still requires optimization.

in DMD, muscle fibers degeneration triggers chronic inflammation, leading to the excessive deposition of extracellular matrix (ECM) proteins. This fibrotic tissue deposition which exacerbates muscle weakness, is a major pathological feature of DMD and significantly contributes to disease progression^13^.

Periostin is a matricellular protein that plays a critical role in tissue repair, inflammation, and fibrosis^14^. It promotes collagen deposition and fibrotic tissue formation in the skeletal muscle^15–17^. Indeed, we recently showed that periostin protein expression is positively correlated to the collagen I and III deposition in the *mdx* mouse diaphragm muscle^15^. Consistently, periostin expression is significantly upregulated in the *mdx* gastrocnemius and diaphragm muscles^18^ and in the *mdx-4cv* diaphragm muscles^19^, 2 mouse models of DMD characterized by degeneration/regeneration cycles.

Alternative periostin isoforms are generated by alternative splicing of exons 16-23 region^20^ with the expression of isoforms containing exon 17 being significantly increased in the diaphragm muscle of muscles of D2.*mdx* mouse compared to the healthy Dba/2J littermate controls^17,21^.

We recently showed that an oligonucleotide antisense strategy, based on periostin exon 17 skipping ameliorated the dystrophic pathology observed in D2.*mdx* mice^21^.

Here, we combined the periostin exon 17 skipping with intramuscular AAV-microdystrophin delivery. The periostin exon 17 skipping significantly improved muscle function, as assessed by running treadmill and forelimb grip strength while it failed to protect the tibialis anterior muscle from repeated eccentric contractions. Notably, microdystrophin protein was significantly expressed by the AAV-MD1 treatment in the tibialis anterior where it protected the muscle from repeated eccentric contractions damage. These data suggest that the combination of these treatments could provide an additive protection and functionality to muscle.

## Methods

### Chronic vivo-PMO-postn Intraperitoneal Injections

All animal procedures were performed in accordance with UK government regulations and were approved by the UK Home Office under Project License P36A9994E. Ethical and operational permission for the in vivo experiments was granted by the Animal Welfare Committee of Royal Holloway University of London. Two-week-old DBA/2J wild-type (WT) and D2.*mdx* mice were randomized by body weight into groups of 4–6 mice. Mice received weekly intraperitoneal injections of 10 mg/kg scramble vivo-PMO (5’-3’: CCTCTTACCTCAGTTACAATTTATA) (vivo-PMO-scr) or periostin exon 17 skipping vivo-PMO (5’-3’: CTTCCGTTTTGATAATAGGCTGAAGACT) (vivo-PMO-postn) from 2 to 12 weeks of age. vivo-PMO-scr and postn (Genetools, USA) were dissolved in 0.9% NaCl solution. The dose was adjusted weekly based on the individual body weight.

### Intramuscular Injection of AAV-MD1

At 6 weeks of age, mice were anesthetized with 2-4% isoflurane and injected with AAV8-Spc5-12-MD1 vector^8^ (low dose: 1e+10 vg/30 µL/muscle; high dose: 4e+10 vg/30 µL/muscle) in both tibialis anterior muscles. Control mice received NaCl (30 µL). Mice were euthanized at 13 weeks of age for tissue collection.

### Forelimb Grip Strength

Forelimb grip strength was measured at 12 weeks using a grip strength meter (Linton Instrumentation, Palgrave Diss, Norfolk, UK). Mice were gently placed on a grid and their tail was pulled until they released their grip. This procedure was repeated five times, with the highest force recorded, normalized to body weight, and averaged. Measurements were performed every two days for consistency.

### Running Treadmill

At 12 weeks, mice were tested for fatigue resistance using a Treadmill Simplex II (Columbus Instrumentation, Columbus, Ohio, USA) set at a 15% incline. After a 5-minute acclimatization, the treadmill speed started at 5 m/min and increased by 0.5 m/min every minute until exhaustion. The distance run before failure was calculated based on the recorded time to fatigue.

### In Situ Tibialis Anterior Muscle Electrophysiology

At 13 weeks, under deep anesthesia (Dolethal 50 mg/kg, Buprenodale 75 µg/kg), tibialis anterior muscle functionality^8,22^ (TREAT-NMD SOP DMD M.2.2.005) was assessed by stimulating the common peroneal nerve. Maximum isometric tetanic force (Po) was recorded at varying stimulation frequencies (10-180 Hz). Specific force (N/cm^2^) was calculated by dividing Po by muscle cross-sectional area. Muscle resistance to eccentric contractions was evaluated by lengthening the muscle after 150 Hz stimulation for 700 ms, followed by a 15% lengthening at 0.75 Lo/s.

### Tissue Collection and Storage

Post-electrophysiology, mice were euthanized by cervical dislocation. Tibialis anterior muscles, diaphragm, and heart were harvested, weighed, and either embedded in OCT medium (VWR, Lutterworth, UK) for histology or snap-frozen for RNA/protein analysis. All samples were stored at -80°C.

### RNA Extraction, cDNA Synthesis, and qPCR

Muscle tissue (30 mg) was homogenized in RLT buffer (QIAGEN, Hilden, Germany) and RNA was extracted using the RNeasy Fibrous Tissue Mini Kit (QIAGEN, Hilden, Germany). RNA concentration and purity were assessed via Nanodrop spectrophotometry. cDNA was synthesized using the QuantiTect Reverse Transcription Kit (QIAGEN, Hilden, Germany). qPCR was performed on the LightCycler480 system (Roche, Mannheim, Germany) with optimized primers (Supplementary Table S1). Gene expression was normalized to Ribosomal Protein Lateral Stalk Subunit P0 (Rplp0)^23^.

### Protein Extraction and Western Blot

Proteins were extracted from frozen muscle tissues using RIPA buffer supplemented with protease and phosphatase inhibitors. Protein concentration was determined using the BCA protein assay. Equal amounts of protein (50 µg) were separated by SDS-PAGE, transferred to nitrocellulose membranes and incubated with primary antibodies: dystrophin (#mannex1011c, DSHB, Iowa City, Iowa, US), periostin (#AF2955; R&D Biosystems, Minneapolis, Minnesota, USA), and α-tubulin (#ab4074, Abcam, Cambridge, UK). Membranes were incubated with secondary antibodies conjugated to 680 or 800 nm fluorophores (Licor, USA) and scanned using the Odyssey CLx system (Licor, Lincoln, Nebraska, US).

### Cryosectioning and Immunofluorescence Staining

Muscle tissues embedded in OCT were cryosectioned (10 µm) at -20°C using a cryostat. For dystrophin immunofluorescence, muscle sections were fixed, incubated with primary antibodies (dystrophin, #mannex1011c, DSHB, Iowa City, Iowa, US; laminin, 1:300, #ab11575, Abcam, Cambridge, UK), followed by secondary antibodies (Alexa 488 and streptavidin-568), thanks to the Mouse on Mouse immunodetection kit (#BMK-2202, Vector Laboratories, Newark, California, USA). Nuclei were stained with DAPI. For Sirius Red staining, muscle sections were incubated in 0.3% Sirius Red solution, dehydrated, and mounted with DPX mounting solution.

### Image Acquisition and Analysis

Immunofluorescence and histological images were captured at 20x magnification using a Nikon NiE Upright microscope (Nikon Corporation, Tokyo, Japan). Dystrophin-positive fibers and Sirius Red-stained areas were quantified using Fiji software. Centro-nucleated fibers were analyzed using the MuscleJ application^24^.

### Statistical Analysis

Data were expressed as mean ± SEM. Outliers were identified using ROUT analysis (Q = 1%) and excluded. Statistical comparisons were performed using one-way or two-way ANOVA followed by Tukey’s post-hoc test. A p-value < 0.05 was considered statistically significant.

## Results

### vivo-PMO-postn improves muscle function of the D2-mdx mouse treated with intramuscular AAV-MD1

10 weeks of intraperitoneal (IP) weekly injections of 10mg/kg of vivo-PMO-periostin (vivo-PMO-postn) or vivo-PMO-scramble (vivo-PMO-scr) were performed in dystrophic D2.*mdx* and littermate control DBA/2J (WT, wild-type) mice from 2 weeks of age. Four weeks after the first injection of vivo-PMOs, 1E+10vg (Low dose) or 4E+10vg (High dose) of AAV-Spc5-12-MD1 were injected in tibialis anterior (TA) muscles. Samples were harvested at week 13 of age. Vivo-PMO treatments were well tolerated by the mice as the weekly body mass measurement showed similar weight gain compared to WT mice (**Figure 1A**).

**Figure 1.**
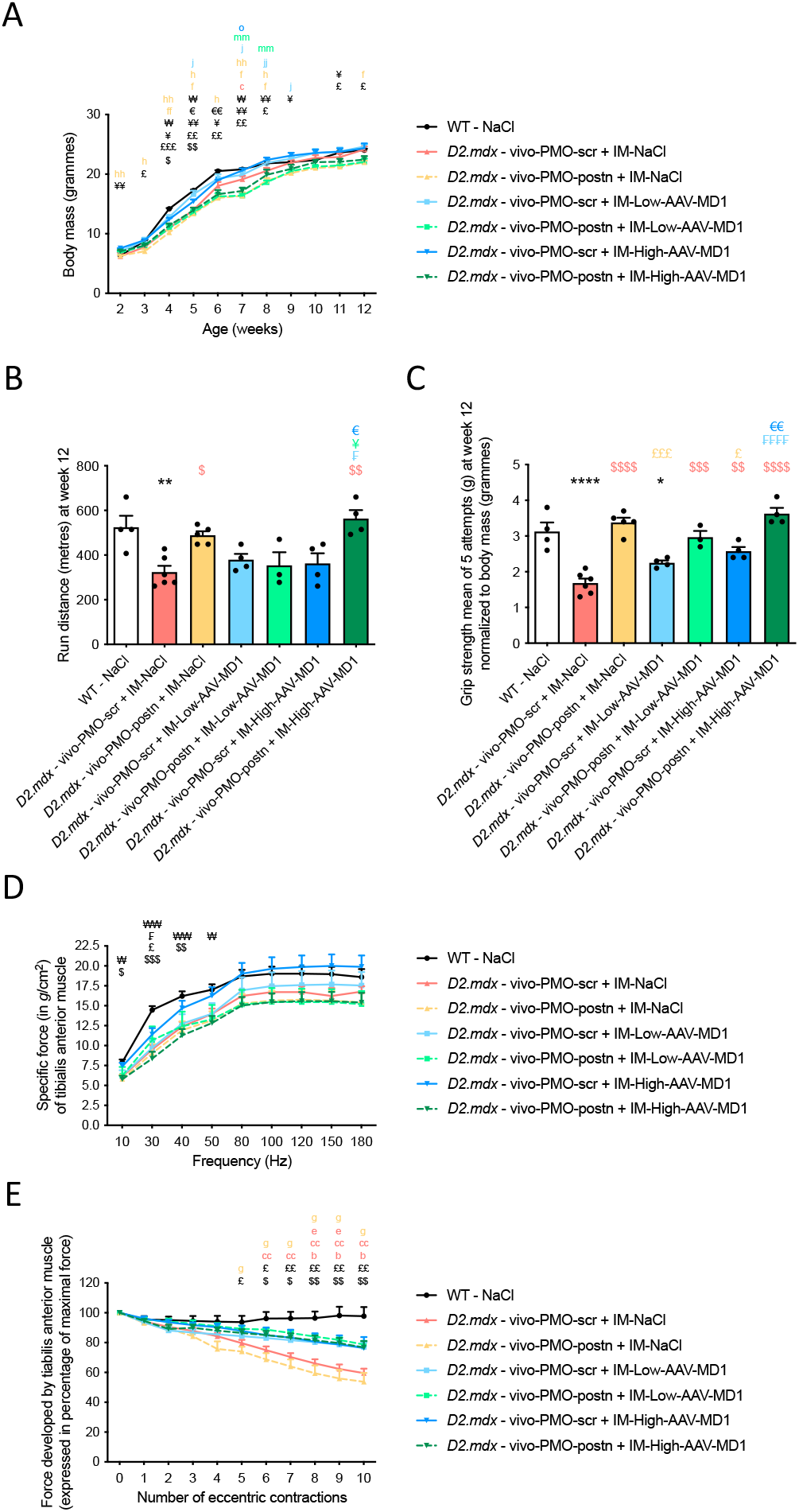
vivo-PMO-postn improves muscle function of the D2-*mdx* mouse treated with intramuscular AAV-MD1. **(A)** Recording of body mass (in grammes) during the experiment duration (from week 2 to week 12 of age), of DBA/2J (WT) and D2.*mdx* mice treated whether with systemic chronic NaCl, vivo-PMO-scr or vivo-PMO-postn injections in combination with an intramuscular NaCl, low or high dose AAV-MD1 single injection, n=3-6 mice/group, ANOVA two-ways followed by Tukey post-hoc comparison test, ^$^p<0.05, ^$$^p<0.01 significant difference between D2.*mdx* – vivo-PMO-scr + IM-NaCl and WT – NaCl, ^£^p<0.05, ^££^p<0.01, ^£££^p<0.001 significant difference between D2.*mdx* – vivo-PMO-postn + IM-NaCl and WT – NaCl, ^¥^p<0.05, ^¥¥^p<0.01, ^¥¥¥^p<0.001 significant difference between D2.*mdx* – vivo-PMO-postn + IM-Low-AAV-MD1 and WT – NaCl, ^€^p<0.05, ^€€^p<0.01 significant difference between D2.*mdx* – vivo-PMO-scr + IM-IM-High-AAV-MD1 and WT – NaCl, ^₩^p<0.05 significant difference between D2.*mdx* – vivo-PMO-postn + IM-IM-High-AAV-MD1 and WT – NaCl, ^c^<p0.05 significant difference between D2.*mdx* – vivo-PMO-scr + IM-NaCl and D2.*mdx* – vivo-PMO-postn + IM-Low-AAV-MD1, ^f^<p0.05, ^ff^p<0.01 significant difference between D2.*mdx* – vivo-PMO-postn + IM-NaCl and D2.*mdx* – vivo-PMO-scr + IM-Low-AAV-MD1, ^h^<p0.05, ^hh^p<0.01 significant difference between D2.*mdx* – vivo-PMO-postn + IM-NaCl and D2.*mdx* – vivo-PMO-scr + IM-High-AAV-MD1, ^j^<p0.05, ^jj^p<0.01 significant difference between D2.*mdx* – vivo-PMO-scr + IM-Low-AAV-MD1 and D2.*mdx* – vivo-PMO-postn + IM-Low-AAV-MD1, ^mm^p<0.01 significant difference between D2.*mdx* – vivo-PMO-postn + IM-Low-AAV-MD1 and D2.*mdx* – vivo-PMO-scr + IM-High-AAV-MD1, °p<0.05 significant difference between D2.*mdx* – vivo-PMO-scr + IM-High-AAV-MD1 and D2.*mdx* – vivo-PMO-postn + IM-High-AAV-MD1. **(B)** Maximal distance ran on a treadmill until fatigue and **(C)** grip strength (mean of 5 repetitions with 30 seconds break between each) (in *g*) developed by forelimbs of by DBA/2J (WT) and D2.*mdx* mice treated whether with systemic chronic NaCl, vivo-PMO-scr or vivo-PMO-postn injections in combination with an intramuscular NaCl, low or high dose AAV-MD1 single injection, at 12 weeks of age, n=3-6 mice/group, ANOVA One-way followed by Tukey post-hoc comparison test, *p<0.05, **p<0.01, ****p<0.0001 significantly different from control WT - NaCl, ^$^p<0.05, ^$$^p<0.01, ^$$$^p<0.001, ^$$$$^p<0.0001 significantly different from D2.*mdx* – vivo-PMO-scr + IM-NaCl, ^£^p<0.05, ^£££^p<0.001 significantly different from D2.*mdx* – vivo-PMO-postn + IM-NaCl, ^₣^p<0.05, ^₣₣₣₣^p<0.0001 significantly different from D2.*mdx* – vivo-PMO-scr + IM-Low-AAV-MD1, ^¥^p<0.05 significantly different from D2.*mdx* – vivo-PMO-postn + IM-Low-AAV-MD1, ^€^p<0.05, ^€€^p<0.01 significantly different from D2.*mdx* – vivo-PMO-scr + IM-High-AAV-MD1. **(C)** Specific force (in *g* force/cm^2^) of tibialis anterior muscle in response to increasing stimulations (10 to 180 Hz) and **(D)** Loss of maximal force (in percentage) during 11 eccentric contractions of tibialis anterior muscle of DBA/2J (WT) and D2.*mdx* mice treated whether with systemic chronic NaCl, vivo-PMO-scr or vivo-PMO-postn injections in combination with an intramuscular NaCl, low or high dose AAV-MD1 single injection, at week 13 of age, n=3-6 mice/group, ANOVA One-way followed by Tukey post-hoc comparison test, ^$^p<0.05, ^$$^p<0.01, ^$$$^p<0.001 significantly different between D2.*mdx* – vivo-PMO-scr + IM-NaCl and control WT – NaCl, ^£^p<0.05, ^££^p<0.01 significantly different between D2.*mdx* – vivo-PMO-postn + IM-NaCl and control WT – NaCl, ^₣^p<0.05 significantly different from D2.*mdx* – vivo-PMO-scr + IM-Low-AAV-MD1 and control WT – NaCl, ^₩^p<0.05, ^₩₩^p<0.01 significant difference between D2.*mdx* – vivo-PMO-postn + IM-IM-High-AAV-MD1 and WT – NaCl, ^b^p<0.05 significant difference between D2.*mdx* – vivo-PMO-scr + IM-Low-AAV-MD1 and D2.*mdx* – vivo-PMO-scr + IM-NaCl, ^cc^p<0.01 significant difference between D2.*mdx* – vivo-PMO-postn + IM-Low-AAV-MD1 and D2.*mdx* – vivo-PMO-scr + IM-NaCl, ^e^p<0.05 significant difference between D2.*mdx* – vivo-PMO-postn + IM-High-AAV-MD1 and D2.*mdx* – vivo-PMO-scr + IM-NaCl, ^g^p<0.05 significant difference between D2.*mdx* – vivo-PMO-postn + IM-Low-AAV-MD1 and D2.*mdx* – vivo-PMO-postn + IM-NaCl.

At week 12 of age, before harvesting samples, treadmill running and forelimb grip strength tests were performed. D2.*mdx* treated with vivo-PMO-scr and IM-NaCl ran less distance (−38%, **p=0.0072) (**Figure 1B**) and provided less forelimb grip strength (−46%, ****p<0.0001) (**Figure 1C**) than the healthy mice. Treatment with vivo-PMO-scr alone restored the running distance (**Figure 1B**) and the grip strength (**Figure 1C**) to the level of the WT mice values. On these functional parameters, the low dose of IM-AAV-MD1 alone did not produced any improvement compared to the untreated D2.*mdx* mice (**Figure 1B-C**). The high dose of IM-AAV-MD1, by itself, did not modify the running distance compared to the untreated D2.*mdx* mice (**Figure 1B**) but significantly increased the forelimb grip strength (+53%, **p=0.004) (**Figure 1C**). The combination of vivo-PMO-postn and low dose of IM-AAV-MD1 has no effect on the running capacity (**Figure 1B**) but significantly increased the grip strength only when comparing with the untreated D2.*mdx* (+115%, ****p<0.0001) (**Figure 1C**). The mice treated with the high dose of IM-AAV-MD1 combined with vivo-PMO-postn ran longer (+55%, *p=0.0161) (**Figure 1B**) and developed more forelimb force (+41%, **p=0.0018) (**Figure 1C**) compared to the mice treated with the high dose of IM-AAV-MD1 alone.

Then, we focused on the TA function using in situ muscle electrophysiology. TA maximal specific force (at 180Hz) was not modified by genotype or treatments (**Figure 1D**). Untreated D2.*mdx* mice were subjected to progressive loss of TA muscle force after eccentric contractions, indicating the fragility of the sarcolemma (**Figure 1E**). While vivo-PMO-postn on its own did not procure any protection against eccentric contraction induced damage, the four groups of mice treated with AAV-MD1, regardless the dose or the combination with vivo-PMO-postn or without, showed the same improvement in membrane resistance, compared to the untreated D2.*mdx* mice (**Figure 1E**).

### vivo-PMO-postn increases MD1 protein expression but does not enhance tibialis anterior muscle histology of the D2-mdx mouse treated with intramuscular AAV-MD1

By western-blot, we observed that microdystrophin protein was restored by both dose of AAV-MD1 injected intramuscularly (**Figure 2A)**. Interestingly, the amount of microdystrophin protein generated by the high dose of AAV-MD1 was further increased by the combination with the vivo-PMO-postn compared to the high-dose-AAV-MD1 alone (fold change x6.78, ****p<0.0001) (**Figure 2C)**. This observation was not made with the low dose of AAV-MD1 (**Figure 2C)**. Dystrophin immunostaining (**Figure 2B**) unraveled a higher percentage of microdystrophin positive fibers with the high dose AAV-MD1 compared to the low dose of AAV-MD1 (+69%, ***p<0.0003), demonstrating a dose effect (**Figure 2D)**. Nevertheless, no improvements were obtained by the combination with the vivo-PMO-postn treatment (**Figure 2D)**.

**Figure 2.**
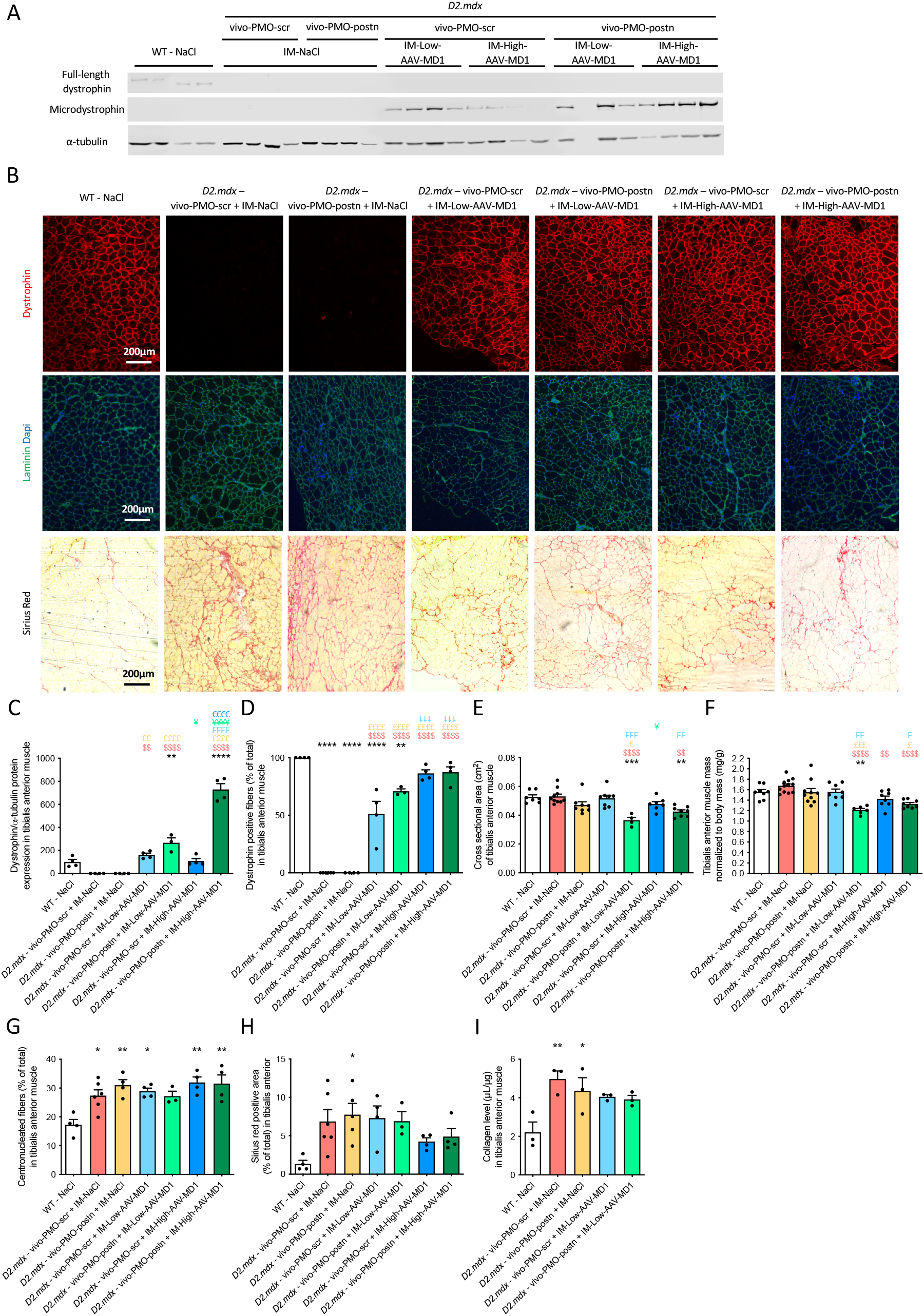
vivo-PMO-postn increases MD1 protein expression but does not enhance tibialis anterior muscle histology of the D2-*mdx* mouse treated with intramuscular AAV-MD1. **(A)** Western-blot of dystrophin, microdystrophin and α-tubulin proteins expression in DBA/2J (WT) and D2.*mdx* mice treated whether with systemic chronic NaCl, vivo-PMO-scr or vivo-PMO-postn injections in combination with an intramuscular NaCl, low or high dose AAV-MD1 single injection, at 13 weeks of age, n=4/group. **(B)** Representative microscopic picture of dystrophin (in red), laminin (in green) and nucleus (Dapi, in blue), and Sirius red stainings, at magnification x100, in 10 µm-tibialis anterior muscle sections of DBA/2J (WT) and D2.*mdx* mice treated whether with systemic chronic NaCl, vivo-PMO-scr or vivo-PMO-postn injections in combination with an intramuscular NaCl, low or high dose AAV-MD1 single injection, at 13 weeks of age. In Sirius red pictures, red labelling indicates collagen I and III staining while muscle fibers appear in light yellow. **(C)** Densitometric analysis of microdystrophin protein signal normalized to α-tubulin, n=3-4 mice/group, results are expressed in percentage of the full-length dystrophin protein expression in WT - NaCl mice, **(D)** quantification of dystrophin positive fibers, expressed in percentage of total number fiber assessed by laminin staining, n=3-4 mice/group, **(E)** cross sectional area (in cm^2^), n=4-7 muscles/group, **(F)** muscle mass (in mg), normalized to the respective animal body mass (in g), n=6-12 muscles/group, **(G)** proportion of centronucleated fibers (in percentage of total), n=3-6 mice/group, **(H)** proportion of Sirius red positive area (in percentage of total muscle area), n=3-6 mice/group, **(I)** collagen level (in µL/µg of muscle tissue), assessed by hydroxyproline assay, n=3 mice/group, in tibialis anterior muscle of DBA/2J (WT) and D2.*mdx* mice treated whether with systemic chronic NaCl, vivo-PMO-scr or vivo-PMO-postn injections in combination with an intramuscular NaCl, low or high dose AAV-MD1 single injection, at 13 weeks of age, *p<0.05, **p<0.01, ***p<0.001, ****p<0.0001 significantly different from control WT - NaCl, ^$$^p<0.01, ^$$$$^p<0.0001 significantly different from D2.*mdx* – vivo-PMO-scr + IM-NaCl, ^£^p<0.05, ^££^p<0.01, ^£££^p<0.001, ^££££^p<0.0001 significantly different from D2.*mdx* – vivo-PMO-postn + IM-NaCl, ^₣^p<0.05, ^₣₣^p<0.01, ^₣₣₣^p<0.001, ^₣₣₣₣^p<0.0001 significantly different from D2.*mdx* – vivo-PMO-scr + IM-Low-AAV-MD1, ^¥^p<0.05, ^¥¥¥¥^p<0.0001 significantly different from D2.*mdx* – vivo-PMO-postn + IM-Low-AAV-MD1, ^€€€^p<0.0001 significantly different from D2.*mdx* – vivo-PMO-scr + IM-High-AAV-MD1.

To assess the histopathological improvement, we performed laminin and nucleus (DAPI) immunofluorescence staining on TA muscle sections (**Figure 2B**). Both the cross-sectional area and the muscle mass of the TA treated with the combination of vivo-PMO-postn and low-dose-AAV-MD1 were reduced compared to muscles treated with low-dose-AAV-MD1 alone (respectively -29%,

***p=0.0002 and -22%, **p=0.0015) (**Figure 2E-F**). On the contrary the percentage of centrally nucleated fibers did not change after treatments and stayed higher in all the D2.*mdx* mice when compared to the WT – NaCl mice (**Figure 2G**).

As periostin is a key pro-fibrotic actor in skeletal muscle, we next explored whether the vivo-PMO-postn treatment affected fibrosis in TA. The Sirius red staining highlights the collagen I and III in red (**Figure 2B**). D2.*mdx* mice presented a higher collagen I and III content compared to the WT – NaCl mice (**Figure 2H**). None of the treatments were able to significantly reduced the TA fibrosis (**Figure 2H**). Similar observations were made when looking at the total collagen content of the TA muscle (**Figure 2I**).

### vivo-PMO-postn is efficient in diaphragm of the D2-mdx mouse treated with intramuscular AAV-MD1 but not in TA or heart

We performed immunoblot and qRT-PCR to study the periostin changes after the PMO. However, no difference in the overall nor 90 kDa main isoform periostin protein expression was observed in muscles treated with either vivo-PMOs (**Figure 3A-C)**. Also, no difference in periostin mRNA was observed for both the full-length isoforms or the isoform excluding exon 17 (**Figure 3D-E**). These results suggest that the treatment did not work in TA muscles.

**Figure 3.**
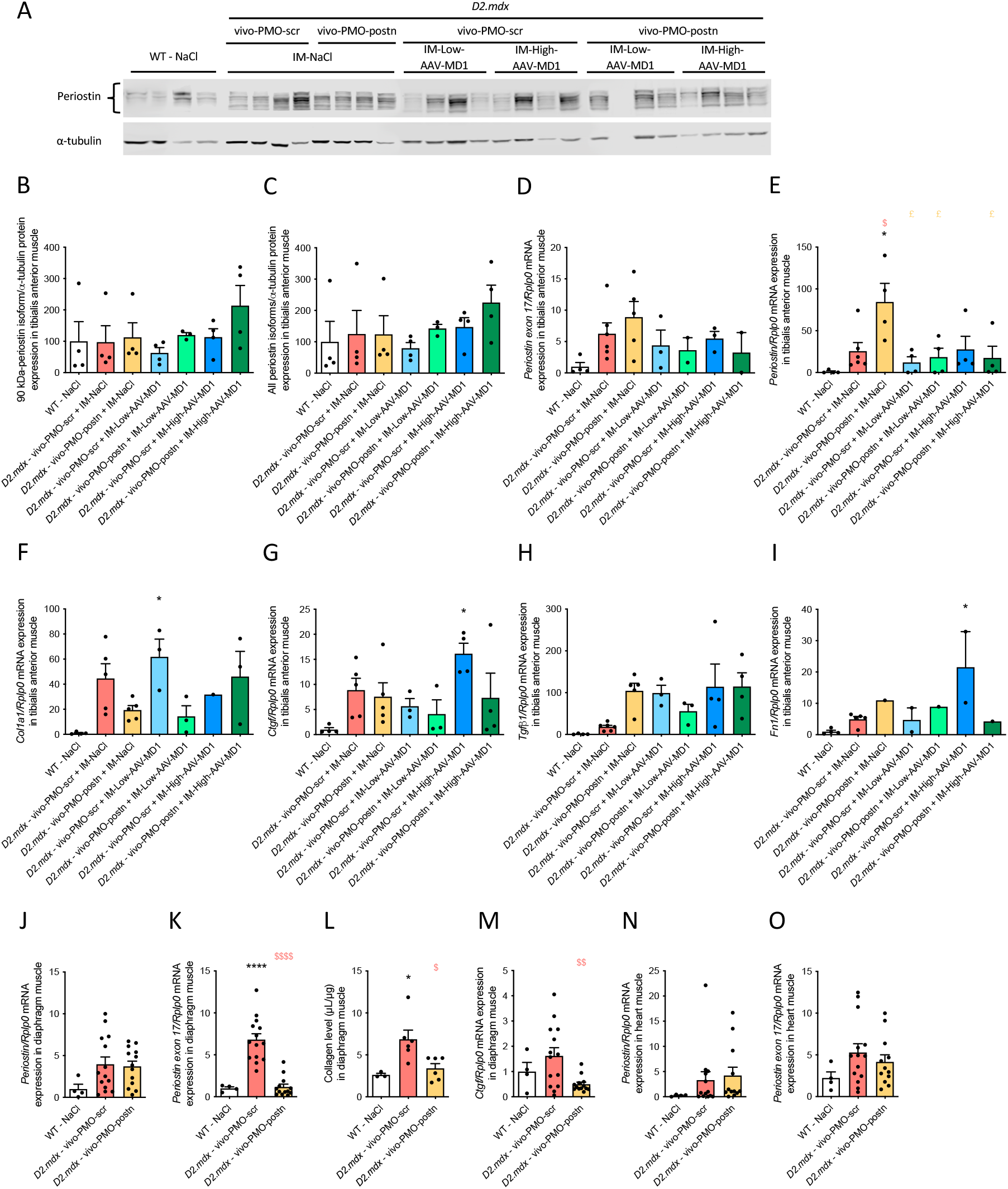
vivo-PMO-postn is efficient in diaphragm and heart muscles of the D2-*mdx* mouse treated with intramuscular AAV-MD1. **(A)** Western-blot of periostin and α-tubulin proteins expression in DBA/2J (WT) and D2.*mdx* mice treated whether with systemic chronic NaCl, vivo-PMO-scr or vivo-PMO-postn injections in combination with an intramuscular NaCl, low or high dose AAV-MD1 single injection, at 13 weeks of age, n=4/group. Densitometric analysis of 90-kDa specific periostin isoform **(B)** and all of them together **(C)**, protein signal normalized to α-tubulin; mRNA expression of *periostin exon 17* **(D)**, total *periostin* **(E)**, Col1a1 **(F)**, Ctgf **(G)**, Tgfb1 **(H)** and Fn1 **(I)** fibrotic genes, assessed by RTqPCR and normalized to *Rplp0* reference gene, in tibialis anterior muscle of DBA/2J (WT) and D2.*mdx* mice treated whether with systemic chronic NaCl, vivo-PMO-scr or vivo-PMO-postn injections in combination with an intramuscular NaCl or AAV-MD1 single injection, at 13 weeks of age, n=4-6 mice/group, ANOVA One-way followed by Tukey post-hoc comparison test, *p<0.05 significantly different from control WT – NaCl, ^$^p<0.05 significantly different from D2.*mdx* – vivo-PMO-scr + IM-NaCl, ^£^p<0.05 significantly different from D2.*mdx* – vivo-PMO-postn + IM-NaCl. Global (*Periostin*) **(J)** and exon-17-containing (*Periostin exon 17*) **(K)** periostin mRNA expression normalized to *Rplp0* reference gene, **(L)** Collagen level (in µL/µg of muscle tissue), assessed by hydroxyproline assay, **(M)** mRNA expression of Ctgf fibrotic genes, assessed by RTqPCR and normalized to *Rplp0* reference gene, in diaphragm muscle of DBA/2J (WT) and D2.*mdx* mice treated whether with systemic chronic vivo-PMO-scr or vivo-PMO-postn injections, at 13 weeks of age, n=4-14 mice/group, ANOVA One-way followed by Tukey post-hoc comparison test, ****p<0.0001 significantly different from control WT - NaCl, ^$^p<0.05, ^$$^p<0.01, ^$$$$^p<0.0001 significantly different from D2.*mdx* – vivo-PMO-scr + IM-NaCl. **(N)** and exon-17-containing (*Periostin exon 17*) **(O)** periostin mRNA expression normalized to *Rplp0* reference gene, in heart muscle of DBA/2J (WT) and D2.*mdx* mice treated whether with systemic chronic vivo-PMO-scr or vivo-PMO-postn injections, at 13 weeks of age, n=4-14 mice/group, ANOVA One-way followed by Tukey post-hoc comparison test, no significant difference.

Next, we analyzed the mRNA of key fibrotic actors like Col1a, Ctgf, Tgf-b1, Fn1 (**Figure 3F-I**). Overall, vivo-PMO-postn and AAV-MD1, by their self or in combination, were not able to reduce mRNA expression of any of these fibrotic markers (**Figure 3F-I**).

Afterwards we analysed the effect of vivo-PMO-postn treatment in other muscles. In agreement with our recent published findings^21^ in diaphragm muscle, vivo-PMO-postn treatment did not change the total periostin mRNA expression (**Figure 3J**) but reduced the periostin exon 17 mRNA expression specifically (fold change x-5.63, ****p<0.0001) (**Figure 3K**). Interestingly, collagen level, assessed by hydroxyproline assay, was reduced by vivo-PMO-postn treatment (−50%, *p=0.024) (**Figure 3L**), consistently with qRT-PCR quantification of the fibrotic marker, Ctgf (−69%, **p=0.008) (**Figure 3M**). Total periostin and exon17 containing mRNA expression were not modified in heart muscle (**Figure 3N-O)**.

## Discussion

In this study, we developed a protocol involving 10 weeks of vivo-PMO-postn systemic injections combined with AAV-MD1 administered intramuscularly in the tibialis anterior (TA) muscle.

Intramuscular injection of AAV-microdystrophin was previously shown to be beneficial in the *mdx* mouse model of DMD generated under the BL10 background^25–28^. Compared to the *mdx* mouse, the D2.*mdx* is characterized by an important muscle fibrosis that reflects more the pathophysiology of human DMD^29,30^. Therefore, it can be considered as a better mouse model to test the combination of microdystrophin restoration and anti-fibrotic periostin exon skipping strategy. We recently showed that the beneficial effects of AAV-MD1 were also observed in D2.*mdx* mice, with a single injection of 4e+12 vg AAV-MD1 at 6 weeks of age in the tail vein^17^. Here, we performed an intramuscular injection of either a low dose (1e+10 vg) or high dose (4e+10g) of AAV-MD1. Both doses were able to restore a significant amount of microdystrophin as detected by western-blot and immunofluorescence. Importantly, if no difference were observed at the total microdystrophin protein expression, we noticed more dystrophin positive fibers with the high dose of AAV-MD1 compared to the low dose, indicating a dose effect. The rationale of using a lower dose of AAV was to avoid saturation of the muscle with MD1, hence the possibility for the vivo-PMO-postn to play an additional role. Consequently, the IM-Low-Dose-AAV-MD1, by itself, and among all the parameters that we analyzed, only offered a significant protection against eccentric contractions damages, compared to the untreated D2.*mdx* mice. Compared with the IM-Low-Dose-AAV-DM1, the IM-High-Dose-AAV-MD1 treatment, on its own, provided some additional benefit as forelimb grip strength was restored to the WT mice level. Overall, this highlights that the AAV8-spc5-12-MD1, with these doses and administration route moderately improves the functionality of D2.*mdx* TA muscle, fulfilling the objective of moderate improvement that can then be further enhanced by vivo-PMO-postn treatment.

Vivo-PMO-postn intraperitoneal injections were applied following the same protocol of our previous study^21^ with 10 mg/kg weekly intraperitoneal injections from week 2 to week 12 of age. Periostin exon 17 skipping was highly efficient in diaphragm muscle, in line with what we observed previously^21^. Based on the known functionalities of the different periostin isoforms^17,21^, this represents a shift from the fibrotic isoform to non-fibrotic isoform with vivo-PMO-postn in the diaphragm muscle. Consistently, the fibrotic marker Ctgf mRNA expression and the collagen content were reduced by the vivo-PMO-postn treatment in the diaphragm muscle. On the other hand, vivo-PMO-postn showed no effect in TA and cardiac muscle. Indeed, neither the periostin mRNA nor the protein expression were modified by the exon-skipping strategy in the TA and in the heart. Consistently, the TA collagens I and III content were not reduced by the vivo-PMO-postn. This difference in efficacy for vivo-PMO-postn treatment could be explained by the different levels of fibrosis which impact TA and heart (low fibrosis), and diaphragm (high fibrosis) muscles of this model^29,30^ which displays a different periostin isoform expression pattern^17,21^. Despite this restricted efficacy to the diaphragm muscle, vivo-PMO-postn treatment rescued the impairment of running capacity and the forelimb grip strength in the D2.*mdx* mice. Further studies would be required to identify other potential muscles and organs that could contribute to the overall muscle function benefits driven by the vivo-PMO-postn, in addition to the improvement in diaphragm function.

In our study we combined for the first time 2 genetic strategies that have previously used independently to improve muscle state and function in the D2.*mdx* mouse model^17,21^. The combination of vivo-PMO-postn with the low dose of AAV-MD1 did not provide any benefits compared to the low dose of AAV-MD1 used alone. Even worse, this combination significantly led to TA muscle cross-sectional area reduction in line with TA atrophy. On the other hand, the combination of vivo-PMO-postn with the high dose of AAV-MD1 significantly increased the AAV-mediated microdystrophin protein expression in the TA. This was not associated with an increase of the proportion of dystrophin positive fibers. Furthermore, we observed that the combination of high dose AAV-MD1 and vivo-PMO-postn, normalized the running capacity as well as the forelimb grip strength, compared to the high dose AAV-MD1 treatment alone. However, the treadmill and grip strength improvements obtained by the vivo-PMO-postn + AAV-MD1 were not significantly different to the ones driven by the vivo-PMO-postn alone, suggesting that these positive effects were mainly driven by the vivo-PMO-postn.

Overall, our results support the rational of combining antifibrotic treatments with strategies restoring dystrophin expression.

## Acknowledgments

This work was funded by a MDUK grant # 22GRO-PG12-0588 awarded to Linda Popplewell.

Dr Alexis Boulinguiez was supported by an AFM-Telethon postdoctoral fellowship and internal pump-prime funding from Royal Holloway, University of London.

We thank Emma Popescu and Claire Gregory for technical help as well as Dr. Penelope Smith and Rob Prouse for logistic support.

## Author Contributions

LP and AM designed the study. JT, AB, NL, JM and AM generated and analyzed the data. AB and AM wrote the first draft of the manuscript. All the authors participated to the writing and validated the submitted version.

