## Supplementary Table S1 for "Periostin exon 17 skipping enhances the efficacy of local AAV-microdystrophin administration in a fibrotic model of Duchenne muscular dystrophy"

<u>Supplementary Table 1:</u> List of primers used for qPCR

| **Primer name** | **Forward 5'-3'** | **Reverse 5'-3'** |
|---|---|---|
| Connective tissue growth factor (*CTGF*) | GTGCACTGCCAAAGATGGTG | CTTTGGAAGGACTCACCGCT |
| Fibronectin 1 (Fn1) | GAGCTATCCATTTCACCTTCAGA | TTGTTCGTAGACACTGGAGAC |
| Periostin (*Postn*) | CCTGTAAGAACTGGTATCAAGGT | CCTTTCATCCCTTCCATTCTCA |
| Periostin exon 17 (*Postn*-17) | ATAACCAAAGTCGTGGAACCAA | CTTCCGTTTTGATAATAGGCTGAA |
| Procollagen 1 (*Col1a1*) | GAAACTTTGCTTCCCAGATGTC | AGACCACGAGGACCAGAA |
| Ribosomal protein lateral stalk subunit P0 (*Rplp0*) | TTATAACCCTGAAGTGCTCGA | CGCTTGTACCCATTGATGATG |
| Transforming Growth Factor (TGFβ1) | CTGCTGACCCCCACTGATAC | GCCCTGTATTCCGTCTCCTT |
